# Functional annotation of a divergent genome using sequence and structure-based homology

**DOI:** 10.1101/2023.08.27.554996

**Authors:** Dennis Svedberg, Rahel R. Winiger, Alexandra Berg, Himanshu Sharma, Christian Tellgren-Roth, Bettina A. Debrunner-Vossbrinck, Charles R. Vossbrinck, Jonas Barandun

**Author notes:** Corresponding author (J.B.). These authors contributed equally to this work (Dennis Svedberg, Rahel R. Winiger, Alexandra Berg).

## Abstract

**Background:** Microsporidia are a large taxon of intracellular pathogens characterized by extraordinarily streamlined genomes with unusually high sequence divergence and many species-specific adaptations. These unique factors pose challenges for traditional genome annotation methods based on sequence homology. As a result, many of the microsporidian genomes sequenced to date contain numerous genes of unknown function. Recent innovations in rapid and accurate structure prediction and comparison, together with the growing amount of data in structural databases, provide new opportunities to assist in the functional annotation of newly sequenced genomes.

**Results:** In this study, we established a workflow that combines sequence and structure-based functional gene annotation approaches employing a ChimeraX plugin, allowing for visual inspection and manual curation. We employed this workflow on a high-quality telomere-to-telomere sequenced tetraploid genome of *Vairimorpha necatrix*. First, the 3080 predicted open reading frames, of which 89 % were confirmed with RNA sequencing data, were used as input. Next, ColabFold was used to create protein structure predictions, followed by a Foldseek search for structural matching to the PDB and AlphaFold databases. The subsequent manual curation, using sequence and structure-based hits, increased the accuracy and quality of the functional genome annotation compared to results using only traditional annotation tools. Our workflow resulted in a comprehensive description of the *V. necatrix* genome, along with a structural summary of the most prevalent protein groups, such as the ricin B lectin family. In addition, and to test our tool, we identified the functions of several previously uncharacterized *Encephalitozoon cuniculi* genes.

**Conclusion:** We provide a new functional annotation tool for divergent organisms and employ it on a newly sequenced, high-quality microsporidian genome to shed light on this uncharacterized intracellular pathogen of Lepidoptera. The addition of a structure-based annotation approach can serve as a valuable template for studying other microsporidian or similarly divergent species.

## Background

Traditional functional gene annotation relies on sequence homology between the studied species and previously characterized genes from other model organisms (*1–5*). However, sequence similarity can be lost over large evolutionary distances (*6, 7*) and, thus, can be very low among highly divergent species (*8–10*). Microsporidia are highly divergent, fungal-like parasites with streamlined and rapidly evolving genomes (*9, 11–13*). As obligate intracellular pathogens, they have been found to infect hosts from almost all animal taxa, including humans (*14, 15*). Besides their medical relevance (*16*), microsporidia infections have a detrimental effect on economically and ecologically important animals such as silkworms and honeybees causing significant economic losses and threatening global food supply (*17–19*). The responsible microsporidian species are among the order of Vairimorpha and Nosema and further analyses of their genomic repertoire and virulence mechanisms are needed to combat microsporidiosis and save important pollinators. Microsporidia develop inside a host cell and are spread to other hosts through an external spore stage. The obligate intracellular nature of microsporidia and the adaptation to this lifestyle has led to the loss of many proteins or sometimes whole biosynthetic pathways. (*11, 20–22*). In addition, microsporidia have shortened not only many of their genes (*9*) but have also reduced intergenic regions to an average of 119 bp in *Encephalitozoon cuniculi* (*5*), leaving them with unusual compacted genomes (*8, 9, 23, 24*). To date, the most extreme case of eukaryotic genome miniaturization is found in *Encephalitozoon intestinalis* at 2.3 Mb with only 1934 densely packed genes (*5*). Despite the reductive evolution, microsporidia have evolved species-specific properties, including a unique and highly specialized polar tube (*25*) for transferring the sporoplasm of the microsporidian to the host cell. The distinctive development of microsporidia, which involves genome reduction, species-specific specialization, and accelerated evolutionary rate, has resulted in significant sequence divergence (*26*).

This divergence observed in microsporidia poses several challenges for traditional sequence-based annotation methods: First, early branching in the fungal kingdom creates a great evolutionary distance to fungal model organisms resulting in diminished sequence similarity (*27*). Second, the accelerated genome evolution, employing gene deletions, mutations, and shortenings as well as enrichments through gene duplications and horizontal gene transfer (from host organisms and bacteria) (*28–30*), shaped a highly divergent clade, not only compared to distantly related organisms but also within the clade itself. Lastly, by optimizing the requirements to infect and thrive in their host (*22, 31*), microsporidia have evolved their own specific set of core genes, which may not exist in other well-studied fungal organisms, such as *Saccharomyces cerevisiae*. In addition, low sequence similarity for universally conserved genes often makes it difficult to find and confirm their homologs in microsporidia.

Unlike primary sequences, protein structures remain more conserved over time (*32, 33*) which is essential to retain their functions (*34*). Proteins with similar functions generally maintain a structural homology (*32, 35*). The gold standard for functional protein annotation is considered to be experimental characterization, including molecular, biochemical, and biophysical analyses. However, the experimental characterization of microsporidian proteins is often not achievable as both culturing and genetic manipulation of microsporidia are challenging. Furthermore, the divergent nature of microsporidian genes, AT-rich genomes, a large fraction of exported disulfide-containing proteins, and codon bias, make it difficult to use typical model organisms such as *Escherichia coli* or *S. cerevisiae* for protein production. Therefore, experimentally verified functional protein annotations lag far behind the amount of sequencing data (*36*). However, recent advances in protein structure prediction provide an improved basis for structure-based functional annotations (*37, 38*). Local, optimized software versions, such as ColabFold (*39*), facilitate creating proteome-wide structure predictions, and Foldseek (*40*), a fast structural aligner, can now be used to search through databases consisting of millions of structures within seconds (*41*).

In this study, we sequenced genomic DNA (gDNA) and total RNA from germinated *Vairimorpha necatrix* (*V. necatrix*) spores, revealing a tetraploid genome with 12 complete chromosomes and 3080 genes. The Benchmarking Universal Single-Copy Orthologs (BUSCO) analysis (*42*) showed a high completeness score of >95%. We used combined structural and sequence-based homology to functionally annotate protein-encoding genes of *V. necatrix*. For this, we developed a ChimeraX plugin to visually inspect every structural annotation match and curate the best hits. Using this approach, we enhanced the prediction of gene function by 10.36% compared to when only relying on sequence-based homology. Further, we found additional, previously unidentified members of the expanded RBL family (*43–45*).

## Results & Discussion

### The genome architecture of *V. necatrix*

We propagated *V. necatrix* in the corn earworm *Helicoverpa zea*, followed by the isolation of highly pure mature spores, which were used for gDNA extraction. The gDNA was sent to the National Genomics Infrastructure Uppsala Genome Center for PacBio *de novo* sequencing and assembly (**Table 1**). The isolated *V. necatrix* spores are tetraploid with 12 completely sequenced and assembled telomere-to-telomere chromosomes (in the following called pseudo-haplotypes 1-4). The assembled pseudo-haplotypes are 15.3, 15.1, 14.8, and 14.7 Mb in size, resulting in a total assembly of 59.9 Mb. The variation in the pseudo-haplotypes’ length is only marginally influenced, for example, by the missing telomers of single chromosomes (max. 20 kb per missing telomer). Overall, the four pseudo-haplotypes share a sequence identity of 96% as assessed by dnadiff (*46*). The differences stem mostly from variations in copy numbers of genes between the chromosomes of the different pseudo-haplotypes. The genome has an overall GC content of 28.3%, and repeated regions make up roughly 50%. To date, assembled microsporidian genomes range from 2.3 Mb (*E. intestinalis*) (*5*) to 51.3 Mb (*Edhazardia aedis*) (*47*), placing *V. necatrix* with an average pseudo-haplotype size of 14.97 Mb among the medium-sized microsporidian genomes. We predicted 3080 open reading frames (ORFs) using BRAKER (*48*), resulting in a coding density of 20.8%. This coding density is on the lower end among microsporidia but is typical for species with a medium-sized to large genome (*49–51*). In comparison, the *E. cuniculi* genome (only 2.9 Mb) has a coding density of 84%. This genome compaction is a result of gene shortening, overlapping genes, and a shortening of intergenic regions (*8, 52*). In the *V. necatrix* genome, however, we only identified three overlapping coding sequences. With a mean intergenic distance of 3606 bp, the intergenic regions are not as significantly shortened as those of *E. cuniculi* (119 bp) (*5*).

**Table 1.** General features of the *V. necatrix* genome.

|  |  |
| --- | --- |
| Haploid genome size (Mb) | 15.3 |
| Ploidy | Tetraploid |
| Repeat content (%) | 50.0 |
| GC content (%) | 28.3 |
| Protein-coding genes | 3080 |
| RNAseq confirmed | 88.7 %, or 2732/3080 genes |
| Gene density (genes/kb) | 0.20 |
| Mean coding length (bp) | 1081 |
| Mean intergenic distance (bp) | 3606 |
| Number of overlapping genes | 3 |
| rDNA genes (16S-23S/5S) | 18 (hap1,3) - 20 (hap2,4) / 13 |
| # of genes with signal peptide | 372 (128 also have a TM) |
| # of genes with transmembrane domains | 382 (128 also have a SP) |
| BUSCO scores of haplotypes | 95.9-96.5% |

To evaluate quality and completeness, we benchmarked the genome against a set of 600 predefined microsporidian-specific genes using BUSCO (*42*), which indicates the presence or absence of highly conserved genes in an assembly. The four pseudo-haplotypes have BUSCO completeness scores from 95.9-96.5% (11 missing and 10 fragmented genes) suggesting a complete genome and accurate ORF prediction (**Table 1**). We used RNA sequencing data to further validate the ORF prediction. For this, RNA was extracted from *V. necatrix* immediately after germination and sent for sequencing. The obtained RNA reads were subsequently aligned to the proteome using STAR (*53*). The ORFs with aligned reads were classified as confirmed (88.7 %, or 2732/3080 genes), and those with no aligned read might either be miss-annotated or not expressed during this early measured time point.

### A ChimeraX plugin for functional genome annotation

Due to microsporidia’s divergent nature and the resulting low sequence identity to proteins in model organisms, many genes’ functions could not be inferred. Similarly, an initial functional annotation of the *V. necatrix* genome with eggNOG (*54*) and based on sequence homology, resulted in 65% hypothetical genes. Previous analyses have shown that structure is often more conserved than sequence, and homologs adopt similar folds despite a very low sequence identity (*32*). Therefore, we conducted a comprehensive structure-based, comparative examination to complement the functional annotation.

First, we used ColabFold to predict protein structures for all identified ORFs in the *V. necatrix* genome and for full proteomes from representative members of the major microsporidian clades (**Figure 1a**). The predicted structures were then matched to the AlphaFold Database (AFDB) and protein database (PDB), using Foldseek in a one-to-all structure-based search (**Figure 1b**). This allowed us to obtain structural similarity scores and top-ranking protein matches. While a structure-based approach can provide complementary functional information on many of the divergent microsporidian proteins, it relies on the quality of the structure prediction and the presence of well-folded domains. Disordered and very small proteins generate only a few structural matches, while ubiquitous domain folds, like short helices, structurally match with many different types of proteins that might be functionally unrelated. Therefore, we concluded that combining a sequence and a structure-based approach, focusing on high-confidence structure predictions, and including a manual curation step for each protein, is best for the functional gene annotation of a divergent organism. To achieve this, we developed a ChimeraX annotator plugin that visually combines the results from structural matches (Foldseek top matches from AFDB, PDB, and in this study folded microsporidian proteomes) with eggNOG (*54*) annotations and the top blast hits (Diamond, (*55*)), while allowing for manual curation (**Supplementary Figure 1, Supplementary Figure 2**). We also displayed transmembrane domain (TMD) and signal-peptide (SP) prediction results in the ChimeraX plugin. We manually curated each protein from *V. necatrix* and updated or complemented the functional annotation. In addition, we used two previous structural studies (*56, 57*) to annotate the ribosomal and proteasomal genes. The high-quality annotation of these proteins can help to improve and correct the functional annotation of other microsporidian organisms. Shortly after finishing our annotation efforts, the automated annotation tool ProtNLM replaced eggNOG as the standard method for gene function prediction. Hence, we compared our annotation results to those of ProtNLM (Benchmarking of our approach). This allowed us to obtain an additional 229 annotations from ProtNLM for gene functions our tool suggested to be uncharacterized or hypothetical.

**Figure 1.**
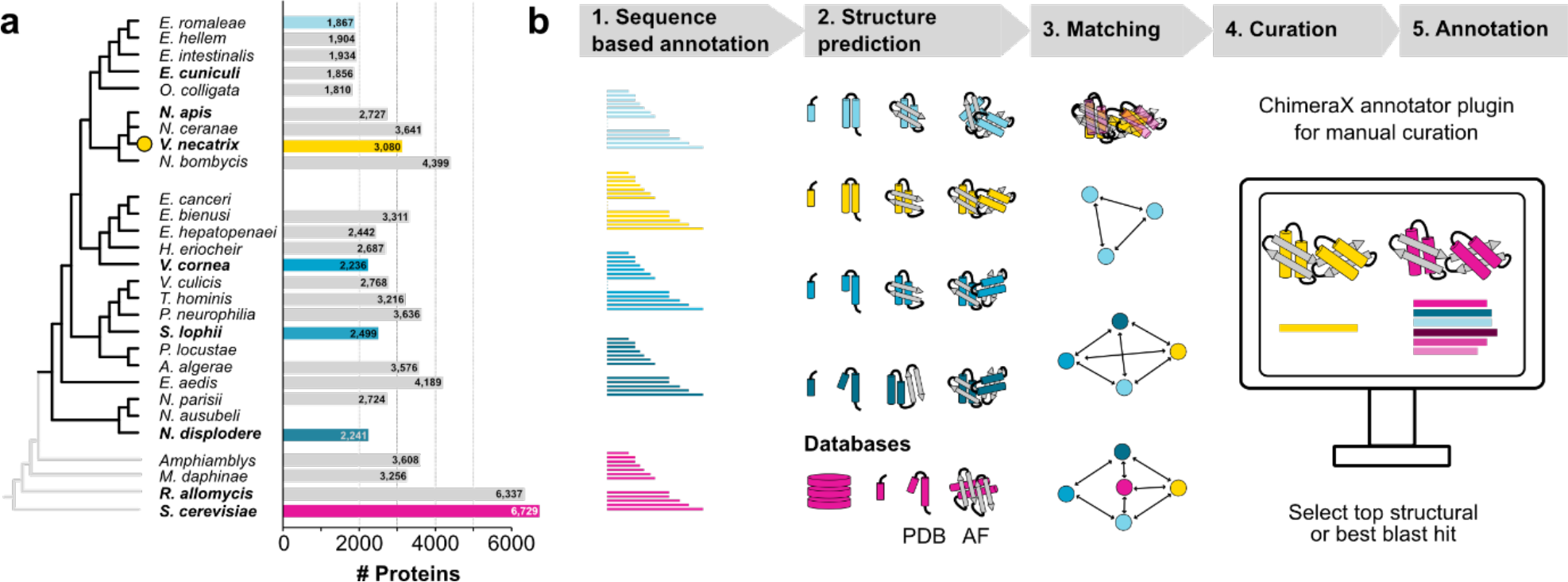
Functional annotation of *V. necatrix* genes using structure-prediction and sequence-based comparative analyses. **a**) A phylogenetic tree based on (24) with 24 microsporidian species, and 3 outgroup species plus *S. cerevisiae* (grey branches). The bar graph shows the number of proteins used and folded for our structural comparison. **b)** Schematic pipeline of our structural homology approach, from protein structure prediction with ColabFold to structural matching using Foldseek, followed by a manual curation step, with our ChimeraX plugin, that includes a comparison of sequence and structure-based hits to achieve a high-quality functional annotation.

The complete manually curated annotations can be found in (**Supplementary Table 1**). The ChimeraX plugin and our annotation database, including all predicted structures, are available as **Supplementary Data File** deposited to Zenodo (10.5281/zenodo.7974739).

### The annotated genome of *V. necatrix*

We employed our ChimeraX annotator plugin on all 3080 predicted proteins that were obtained from ORFs of the *de novo* assembled *V. necatrix* genome. We functionally annotated 1932 proteins in total using combined information from sequence, predicted structural, and available experimental data (**Figure 2a, Final curated**). Compared to eggNOG and ProtNLM, we were able to annotate an additional 319 genes in the *V. necatrix* genome, excluding the information from experimentally verified proteins. Experimental data allowed us to unambiguously identify proteasomal (*57*) and ribosomal (*56*) genes (**Figure 2a, Experimental)**.

**Figure 2.**
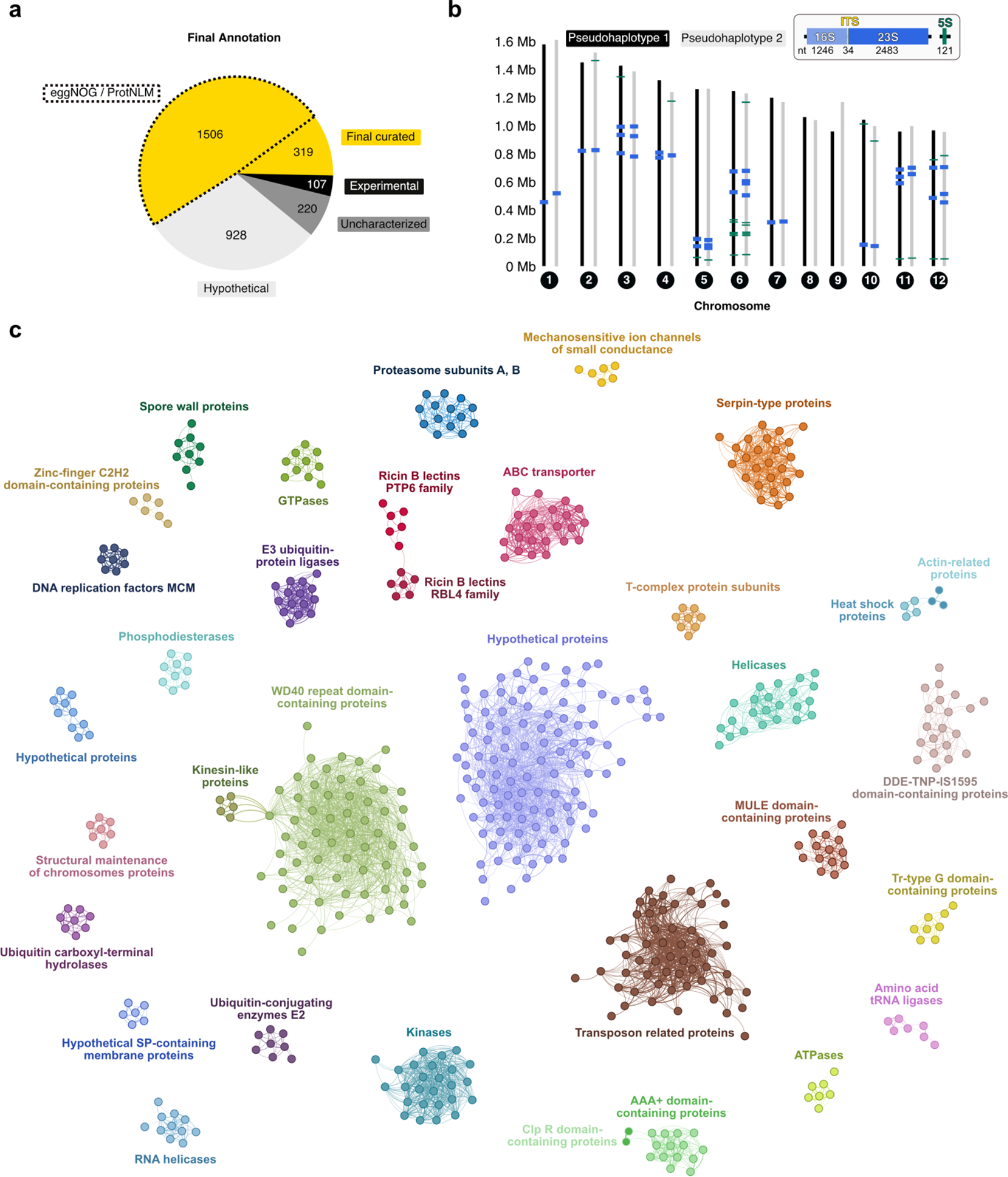
The annotated genome of *V. necatrix*. **(a)** Pie chart summarizing the functional annotation output using a combination of sequence and structure-based hits and experimental data. Compared to ProtNLM or eggNOG (yellow, marked by black dashed lines), our complementary approach improved the genome annotation by an additional 319 final curated gene functions, here shown in yellow. Further, 107 experimentally solved protein structures (black) from PDB are listed as structural matches. 220 genes that have homologs in other microsporidia, but are of unknown function, are presented in dark grey. Light grey represents 928 hypothetical *V. necatrix* genes that have no matches to the known genes of other microsporidia. **(b)** Approximate localization of the rDNA genes 16S/23S (blue) and 5S (green) on the 12 chromosomes of the two predominant haplotypes 1 (black) and 2 (grey). The insert depicts one rDNA in shades of blue (light blue for the 16S, dark blue for the 23S) and one 5S gene in green. The internal transcribed spacer (ITS) is shown in yellow. **(c)** Structure-based network of highly enriched protein-fold families encoded by our *V. necatrix* genome. Alphafold-predicted protein models were analyzed for structural relatedness in a Foldseek all-against-all search. The structural similarity is represented by the TM score which is used as a measure for the protein network graph generated in Gephi. Each node represents a protein colored according to its fold family. Proteins with inverted surrounding and filling color compared to the main cluster have an additional common domain besides the one unifying the main cluster i.e., Clp R domain-containing proteins and actin(-like) proteins. Connecting lines indicate structural relation of proteins and thicker lines indicate greater structural similarity. PTP6, polar tube protein 6; RBL, ricin B lectin; MCM, minichromosome maintenance; Serpin-type protein, serine-inhibitor type protein; MULE domain, Mutator-like elements domain; Tr-type G domain, translation-type guanosine-binding domain; SP, signal peptide; Clp R domain, caseinolytic protease repeat domain; AAA+, ATPases associated with diverse cellular activities.

Some of these genes were either not predicted or falsely predicted when using a sequence-based approach. Of the annotated genes, 92% were confirmed by RNA reads. A total of 1148 *V. necatrix* genes could not be functionally annotated (**Figure 2a**) using either traditional or structural annotation tools, as homology hits were missing or of low confidence. Of those, hypothetical genes that are conserved in several microsporidian species were termed “uncharacterized”, whereas others that are only conserved within the order Nosematida were called “hypothetical”. RNA reads covered >87% of the hypothetical genes suggesting that most hypothetical genes are expressed and not a result of an overestimated number of protein-coding regions predicted by BRAKER.

The rRNA sequence, validated with the ribosome structure, allowed us to map the rDNA genes with high confidence (**Figure 2b)**. The pseudo-haplotypes contain 18 (pseudo-haplotype 1 and 3) or 20 (pseudo-haplotype 2 and 4) full rDNA loci, which are not clustering as repeats or localizing to subtelomeric regions as observed in *E. cuniculi* (*58*). The rDNA loci are distributed among all chromosomes except for chromosomes 8 and 9. About half of the 12 to 14 copies of the 5S gene localize to chromosome 6, while the additional copies are distributed on other chromosomes and generally closer to the chromosome ends (**Figure 2b**).

To obtain a global view of the most abundant protein folds in *V. necatrix*, we performed an all-to-all Foldseek search and visualized structurally related proteins with a network graph (**Figure 2c)**. The most enriched protein fold in *V. necatrix* is represented by WD40 repeat domain-containing proteins followed by transposon-related proteins. While WD40 repeat domains are known as one of the most plentiful domain families in eukaryotes and are involved in protein-protein interactions (*59, 60*), the abundance of transposon-related elements in microsporidia can vary with the size of the genome. Overall, gene-sparse microsporidian genomes range from 12 to 50 Mb in size, and their non-coding regions are predominantly found to be transposable elements (*61*). In our 15 Mb *V. necatrix* genome, we annotated around 109 retrotransposable elements that are involved in genetic mobility and genomic plasticity (**Figure 2c**). In contrast, in the gene-dense genome of *E. cuniculi* (2.9 Mb), no such elements or RNA-dependent reverse transcriptases were identified, apart from the telomerase reverse transcriptase (*8*). Additionally, we identified a Dicer-like protein (VNE69_01137) and an Argonaute protein (VNE69_01023), which belong to the RNAi machinery. A functional RNAi pathway correlates with a higher proportion of transposable elements and larger genome sizes (*61*) which might explain the high number of transposable elements found in *V. necatrix*.

Apart from eukaryotic conserved protein families (e.g., ABC transporter, kinases, AAA+ ATPases, ABC transporters), *V. necatrix* harbors a large amount of Serine protease inhibitors (Serpins), ricin B lectin (RBL)-like proteins (discussed below), and spore wall proteins (**Figure 2c**). To date, Serpins were exclusively found in Nosema (*62*), a genus infecting insects. One of the defense mechanisms of insects against pathogens is hemolymph melanization, which relies on the serine protease-mediated prophenoloxidase activation cascade. This process results in the inactivation of pathogens due to the deposition of melanin onto the invaders. Microsporidian Serpins were suggested to be secreted during host invasion to inhibit the prophenoloxidase activating proteinase, thereby interfering with the host’s innate immune response (*21, 63*). This melanization pathway is conserved in Lepidoptera (*64, 65*), the host of *V. necatrix* (58), providing a potential reason for the enriched repertoire of Serpins we identified in this study. Furthermore, the outermost layer of the mature microsporidian spore was shown to include many SWPs (*66*). Since the spore wall is thought to be the first and most direct contact point with the environment and the host cell, the SWPs have potentially crucial roles in signaling, adherence, or enzymatic interactions (*67*). Further studies are required to analyze the importance of these protein families for parasite adherence, invasion and host immune evasion mechanisms.

### Enhancing automated annotations through structural homology searches followed by manual curation

To benchmark our approach, we compared our method’s final gene function annotations to the results from the ProtNLM automated annotation (**Figure 3a**). In a second step, we employed our workflow on the uncharacterized genes from *E. cuniculi* retrieved from UniProt (accessed October 2022) (*8*), to test if we can improve the annotations that were recently updated with ProtNLM (*68*). When comparing our annotations with the assignments of ProtNLM, 42% of the gene function predictions had the same name or were homologous, and 33% were different at first glance and were further looked at in more detail below. Both approaches failed to annotate 22% of all gene functions, while 3% could be assigned using solved protein structures (*56, 57*). The 42% identical annotations, included cases where one approach predicted a gene function while the other identified a protein domain typically fulfilling this gene function, or vice versa. These identical predictions gave a higher confidence in the assigned gene function. Among the 1009 differing annotations were 639 predicted gene functions made by us, for which ProtNLM provided low-confidence predictions with a model score below 0.2, an exclusion threshold used for UniProt annotations (https://www.uniprot.org/help/ProtNLM). Further 370 different hits contained 229 uncharacterized or hypothetical proteins according to our approach. For these, we assigned the predicted domain characterization and/or gene function from ProtNLM based on their E-value > 0.2. Additionally, 14 microsporidia-specific genes were predicted by our approach with high confidence and were not recognized by ProtNLM. The remaining 126 seemingly differing annotations required a closer look. Some proteins were assigned with a different (domain) function i.e., transposable elements (endonuclease vs. integrase), by us compared to ProtNLM. However, the protein might as well harbor both domains or fulfill both functions but biochemical analysis is necessary for confirmation. Further disparate annotations included non-informative predictions by ProtNLM such as DUF (domain of unknown function), phage protein, or WD40-repeat domain-containing proteins. For these cases, our manual curation step, which allowed us to visualize the protein of interest, increased the confidence in our functional prediction of the proteins. Further, up to 4% (121 genes) within the non-identical predictions include potential miss-annotations by ProtNLM (**Figure 3a**). Of these, 4%, 61 gene functions are related to other obligate intracellular pathogens such as apicomplexans. The genes include surface and secretory proteins that aid in the parasitic lifestyle and are associated with the invasion into a host cell, formation of a parasitophorous vacuole, and replication. Since roughly one-third of these genes have a prediction model value above 0.2 in ProtNLM, it is likely that microsporidia share certain protein features with other intracellular parasites (*69, 70*). However, instead of automatically carrying over the exact annotation i.e., “oocyst capsule protein”, we suggest annotating these as “oocyst capsule protein-like”.

**Figure 3.**
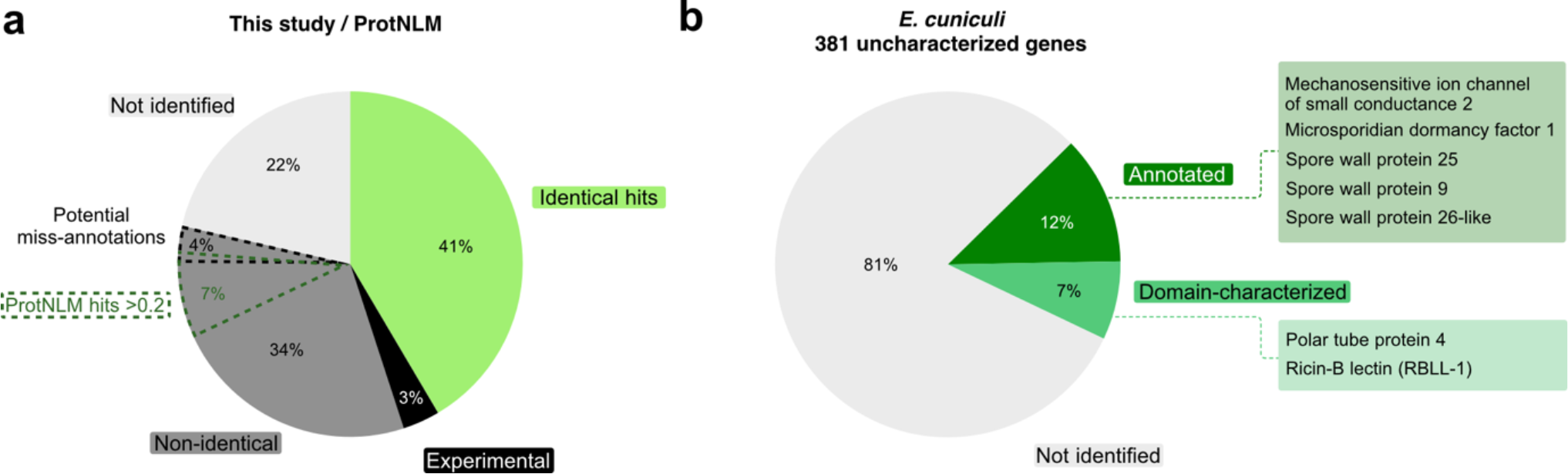
Complementation of structure and sequence-based functional annotation enriches the total number of matches and improves the annotation of microsporidia-specific genes. **(a)** To assess the annotation efficiency of our combined structure and sequence-based homology approach, we counted the amount of identical (green), non-identical (dark grey), not identified (light grey) and experimentally determined (black) functional gene predictions between our ChimeraX tool and ProtNLM. Additionally, we display the relative number of potential miss-annotations (dark grey with black dashed line) predicted by ProtNLM and the percentage of ProtNLM gene function predictions with a model value above 0.2 (dark green dashed line) that we transferred to genes our approach suggested to be uncharacterized or hypothetical. (**b)** Employing our approach, we functionally annotated 12% (dark green) and characterized the domain of 7% (green) of the 381 uncharacterized *E. cuniculi* proteins. RBLL-1, ricin-B lectin-like 1.

We next tested our structural homology approach on the 381 uncharacterized proteins from *E*. *cuniculi* (strain GB-M1) (*8*), for which the current functional prediction is sequence-based and was recently updated with ProtNLM annotations on UniProt (*68*). By manually curating every protein, we could functionally annotate 46 proteins, and characterize domains in 26 proteins **(Figure 3b**, **Supplementary Table 2)**. Our approach showed a clear advantage for microsporidia-specific genes that encode polar tube proteins and spore wall proteins and for proteins characterized via experimental structural analyses. We identified three microsporidia-specific spore wall proteins (Q8SVI9, Spore wall protein 25; Q8SV25, Spore wall protein 9; Q8SVK8, Spore wall protein 26-like), two RBLs (Q8SUK2, ricin B lectin (Polar tube protein 4); Q8SUY7, ricin B lectin-like protein 1 (RBLL-1)) and the microsporidian dormancy factor 1, MDF1 (Q8SWQ4). Further, we annotated the gene coding for the mechanosensitive ion channel of small conductance 2 (Q8STV6), of which only a single copy exists in every microsporidian genome sequenced to date (*30*). None of these assigned protein functions or domains were identified by ProtNLM, suggesting that structural homology is an important complementary approach to predicting protein functions and characteristics in divergent organisms.

### For many divergent microsporidian proteins, structures are more conserved than sequences

Using structural homology matching, we identified several proteins that were not identified previously through sequence similarity alone. For example, we attempted to retrieve the eleven genes that were not identified during the BUSCO search, which is purely sequence-based (**Table 1**, **Figure 4a)**. Since large differences in the gene content can occur within higher taxonomic levels, it is necessary to use a specific BUSCO data set for the species of interest (*71*). For microsporidia, the set of 600 reference genes to determine the BUSCO score stems from the Encephalitozoon genus (**Figure 4a**). We thus folded the corresponding *E*. *cuniculi* proteins using ColabFold and performed structure-based matching with Foldseek to the *V*. *necatrix* proteins encoded by the predicted ORFs. With high confidence, four out of eleven missing genes were identified, increasing the BUSCO score slightly. The identified proteins displayed a high TM score but a low sequence identity. The additionally matched proteins were the Endoplasmic reticulum membrane-associated oxidoreductin (ERO1), Mitochondrial import inner membrane translocase subunit TIM50, High-mobility group protein, and Ribosomal protein eS10 (**Figure 4a**, and **4b**). An additional protein (RING-type E3 ubiquitin transferase) that was identified had a low TM score, potentially due to disordered regions and high flexibility linkers. For the other six proteins, no clear best hit could be retrieved.

**Figure 4.**
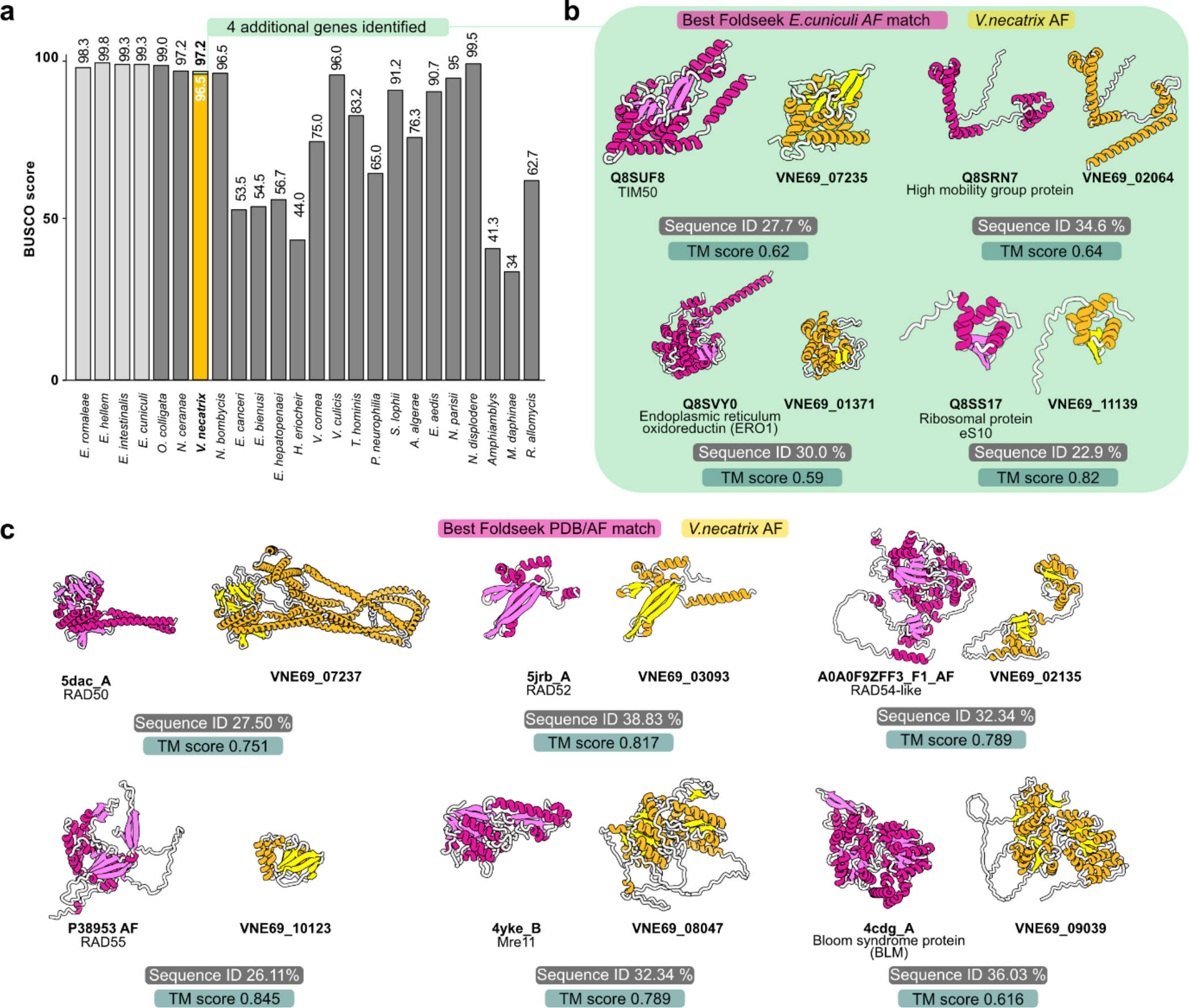
Examples of high-confidence structure-based hits with low sequence identity. (**a**) BUSCO scores of a selection of microsporidian genomes compared to the score of *V. necatrix*. The genus Encephalitozoon is colored light grey. The *V. necatrix* BUSCO score bar is colored yellow with an extension in green representing the four additional genes identified using Foldseek. (**b**) AlphaFold structures of *E. cuniculi* (magenta) and *V. necatrix* (gold) proteins corresponding to the four microsporidia BUSCO genes. These four genes were exclusively identified via structural matching due to their low protein sequence identity. (**c)** A subset of HRR proteins identified in *V. necatrix*. Complementing the traditional sequence-based approach with our structural homology searches allowed for the identification of most proteins involved in the HRR. Displayed proteins have a high TM score above 0.5 and a sequence identity below 40%. Sequence similarity was calculated with Clustal (v2.1), and TM scores were generated using TM-align (https://zhanggroup.org/TM-align/). TM score was normalized according to the length of the reference protein. Reference protein IDs containing “AF” refer to AlphaFold-predicted structures. Gold: Identified microsporidian proteins; magenta: Homologs; AF, AlphaFold; PDB, Protein Data Bank.

Additionally, we used structural homology searches to help identify proteins involved in DNA repair pathways that otherwise might be overlooked by or ambiguous to sequence-based tools. While our study supports previous findings (*72*) that the non-homologous end joining pathway appears to be absent or highly reduced in microsporidia, we found most of the key genes listed in the KEGG pathway database, involved in the homologous recombination repair (HRR) system (**Figure 4c**). However, Rad57 could not be unambiguously identified. In addition, neither yXrs2 nor hNbs1, belonging to the MRX respectively the MRN complex, could be detected, which suggests that they are either absent or too sequence-divergent to produce a reasonable structural model. Overall, this may indicate that *V. necatrix* uses a minimized version of HRR for DNA repair.

### Structure-based identification and classification of the expanded RBL family in microsporidia

Large expanded gene families in obligate intracellular pathogens are postulated to have an important role in host-pathogen interactions (*73*). Among microsporidia, leucine-rich repeats are significantly enriched in the order Nematocida (*74*), while Serpins (*62, 63, 74*) and RBLs are highly abundant in Nosematida (*43–45, 74*). For example, 52 RBL proteins were identified in the honey-bee pathogen *Nosema bombycis* (*45*) where they were shown to enhance spore adhesion and host-cell invasion (*43*). However, the authors predict that 22 of these proteins lost their RBL domain due to extreme sequence divergence in microsporidia (*45*). An alternate explanation might be that sequence-based methods are insufficient to identify the RBL domain or that previously, proteins were erroneously annotated as RBL-domain-containing proteins. To test this hypothesis, and to unambiguously detect RBLs present in microsporidia we complemented the existing sequence-based search with our approach. For this, we focused on RBLs in Nosematida to demonstrate our structure-based workflow for finding homologs with low sequence identity.

We identified a total of 74 RBLs of which 22 were found in *V. necatrix* and several previously not identified in other species. We clustered them into 13 different RBL clades (**Figure 5a**), of which four contain previously characterized proteins. These four include the PTP4, PTP5, PTP6, and RBLL-1 clades (**Figure 5a)**. Proteins from these clades all localize to different parts of the microsporidian polar tube (*75–78*) or the spore wall (*44*). PTP4 and PTP6 were shown to mediate host cell binding, while RBLL-1 from *E. cuniculi*, was shown to interact with the polar tube and spore wall (*44*). The remaining nine uncharacterized RBL groups in our cladogram were termed RBL1 through 9. The clades RBL1 and RBL2, similar to PTP4 and PTP5, are conserved among microsporidia, as all species (except for *O. colligata* in the PTP4 and PTP5 clades (*79*)) are represented with one gene each. In contrast, members of the clades PTP6 and RBL4 were only identified in *V. ceranae* and *V. necatrix*. The RBLL-1 clade is represented exclusively by Encephalitozoon species, with *E. cuniculi* harboring the most RBL proteins (Q8SUK1 through 4). In the PTP4, PTP5, PTP6, RBL4, and RBLL-1 groups, almost all corresponding genes form clusters in the respective microsporidian genomes. *ptp4* and *ptp5* are always adjacent in all the genomes analyzed in this study, a finding previously reported for other microsporidian species (*76*). Most of the *ptp6* genes are localized adjacent to *rbl4* genes in our *V. necatrix* genome (**Supplementary Table 1**), and the four *rbll-1* genes in *E. cuniculi* form a gene cluster as well. These lineage-specific enrichments of *rbl* genes could result from gene duplication events in response to the host immune system during infection and the subsequent evolutionary pressure on microsporidia to re-optimize host-cell attachment. Alternatively, their genomic closeness could indicate functional or physical interaction.

**Figure 5.**
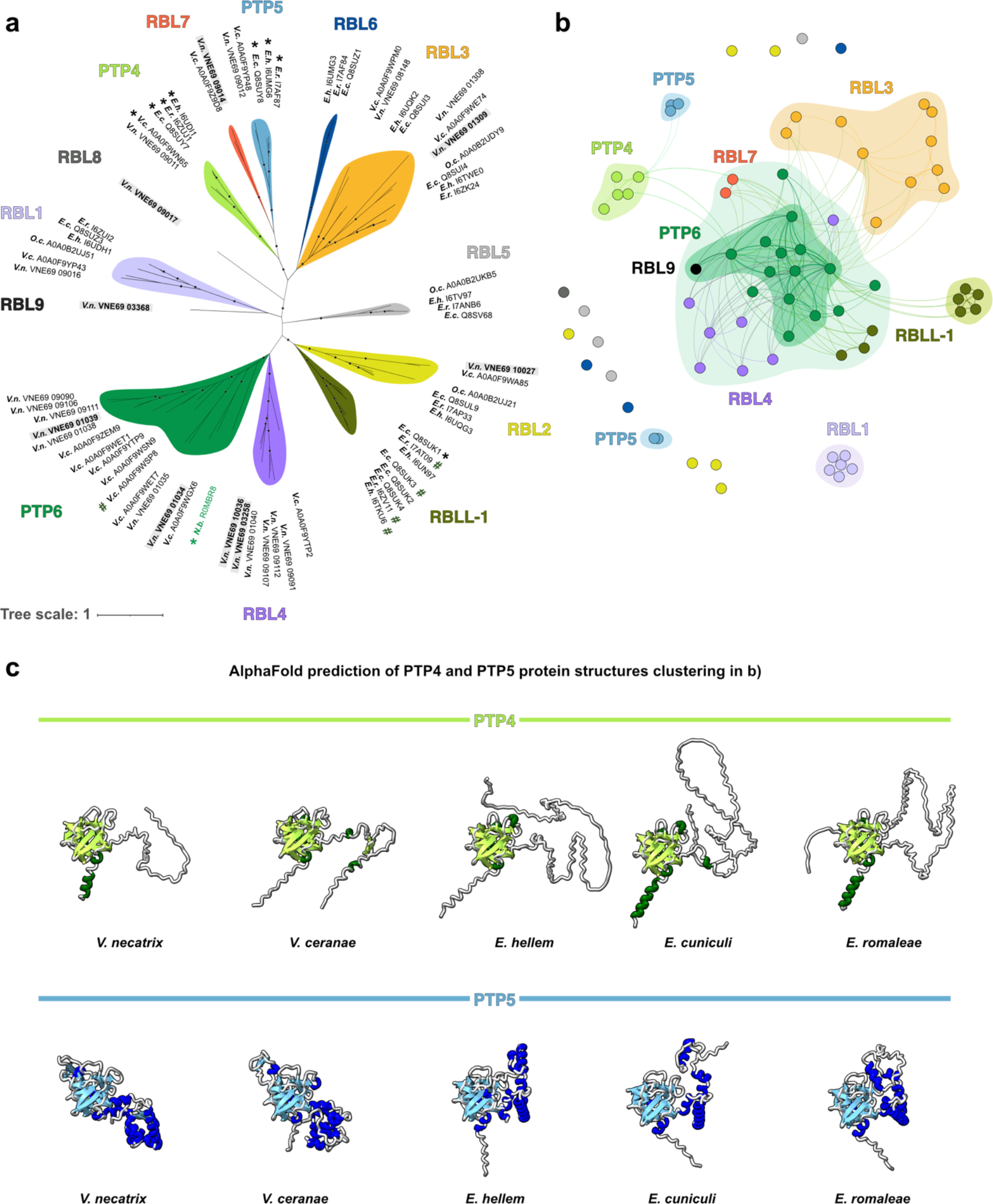
Structure-based identification and classification of the abundant RBL protein family. **a**) Cladogram of Nosematida RBLs named based on available experimental data (PTP4, PTP5, PTP6, RBLL-1) and otherwise termed RBL1 through RBL9. Branches marked with stars indicate a bootstrap value >70. Protein IDs with asterisks indicate existing publications on the respective gene, hashtag marks indicate previously identified orthologs to NbPTP6 (77), and proteins in bold with a light grey background indicate that corresponding genes are expressed during germination. **b**) Structure-based network of RBLs color-coded according to their clade in a). Each node represents one RBL protein, connecting lines indicate the degree of structural relatedness, and surrounding shapes in brighter shades mark structural clusters. Protein folds of all RBLs identified in a) were predicted with AlphaFold and RBLs were clustered according to structural similarity based on their TM score using Gephi (82). **c**) AlphaFold-predicted protein structures for the PTP4s and PTP5s comparing tertiary structures between the two protein families and the microsporidian families. *E*.*c*., *Encephalitozoon cuniculi*; *E*.*h*., *Encephalitozoon hellem*; *E*.*r. Encephalitozoon romaleae*; *N*.*b*., *Nosema bombycis; O*.*c*., *Ordospora colligata*; *V*.*n*., *Vairimorpha necatrix*; *V*.*c*., *Vairimorpha ceranae*; RBL, ricin B lectin; RBLL, ricin B lectin-like; PTP, polar tube protein.

Next, to analyze the structural relationship among Nosematida RBLs, we generated a structure-based protein network using the TM score as a measure of structural similarity (**Figure 5b, Supplementary Figure 3**). We found that most sequence-based RBL groups in the cladogram (**Figure 5a**) also cluster structurally, suggesting conserved function within the respective RBL clades and across species of the Nosematida order. Exceptions are the PTP5, RBLL-1, and RBL4 members, which form two structural groups each. However, most RBL proteins show structural similarity to the PTP6 clade members which form a center in this network. A total of 14 RBLs form outliers that are completely unrelated to any other RBL group; these include all members of the RBL2, RBL5, RBL6, and RBL8 clusters. Additionally, the RBL1s have no connection to neighboring RBL classes but form a joint cluster with all clade members identified in **Figure 5a**.

Since both PTP4 and PTP5 are unique to microsporidia and form part of their infection apparatus, we were interested in their structural similarities and differences, based on the AlphaFold predictions (**Figure 5c**). The structural network shows PTP4s clustering together, while PTP5s are divided on a genus level (**Supplementary Figure 3**): *V. necatrix* and *Vairimorpha ceranae* PTP5 seem structurally less related to the Encephalitozoon homologs, possibly due to an additional β-sheet pair, incorporated in the RBL domain (**Figure 5c, lower panel**). Apart from that, both polar tube proteins, PTP4 and PTP5, share a highly conserved and structurally nearly identical RBL domain but differ in their N- and C-termini. The N-terminally predicted signal peptide varies in secondary structure. The C-terminal regions of all PTP4 are predicted to be disordered (**Figure 5c, upper panel**) while those in PTP5 are more structured (**Figure 5c, lower panel**). A disordered loop is thought to be advantageous when facing host-immune response as it is more flexible in adopting various conformations (*73*) and to date only PTP4, but not PTP5 has been shown to mediate host-cell binding (*75*).

Structural homology searches allowed us to identify new members of the large RBL protein family in the Nosematida. We also showed that PTP4-6 are members of the RBL family, which may contain additional, yet uncharacterized proteins that form part of the unique microsporidian infection apparatus. A recent study identified multiple RBL proteins as interaction partners of *V. necatrix* PTP3 (*80*), one of the main components of the polar tube (*81*). Our findings of the close relationship between PTP4-6 and RBL proteins, and the interaction of various RBL proteins with the microsporidian polar tube indicate an important role of RBLs in microsporidian host-invasion and incentivize further experimental research on this enriched protein family in Nosematida.

## Conclusion

The functional annotation of proteins is a critical step for understanding the biology of organisms. Even though automated annotations are essential to whole genome/proteome projects, they traditionally rely on sequence identity, orthology searches, and protein name predictions based on the amino acid sequence. This poses three major problems: First, sequence similarity searches can fail to result in significant matches if the sequence is too divergent from the ones present in databases. This is often the case when analyzing understudied species like microsporidia or newly emerging pathogens. Second, up to date, low sequence-identity blast hits against *S. cerevisiae* and other model organisms led to functional annotation of microsporidian genes that are neither in accordance with the structural hits identified by Foldseek nor with the ribosomal and proteasomal genes revealed through structural studies (*56, 57*). Third, any previous annotation error is likely to be propagated across species. Thus, for divergent species with low sequence identity like microsporidia, sequence-based annotations are not sufficient. However, since the structure and the biological role of a protein are connected, protein function can be inferred using structural homology searches. We developed a functional annotation workflow that allowed us to manually curate sequence and structure-based matches and to select the best hit based on sequence identity and TM score. We used this annotation workflow on our newly sequenced, high-quality genome of *V. necatrix*, a microsporidian species poorly characterized up to this point.

The implementation of structural homology searches and the manual curation step, that our plugin offers, allows us to identify potential miss-annotations and may thus prevent their automatic transfer in the future. Further, it is possible to filter out proteins that are exclusively present in invertebrates and are thus most likely contaminants. Our pipeline, complemented with protNLM, allowed us to functionally annotate 1932 out of 3080 predicted genes, including an additional 319 hits compared to annotations using traditional sequence-based approaches only. The complementary information from sequence and structure further allowed us to characterize 19% (72 proteins out of 381) of the *E. cuniculi* proteins or protein domains that were previously annotated as “hypothetical” or “uncharacterized”. Further, using structural homology searches, we have identified previously unknown RBL family members in the order Nosematida and shown that PTP4, PTP5, and PTP6 are part of the RBL family. Structural information gives a first hint of the putative function of a protein, its structural appearance, and potential interaction partners and may thus provide guidelines for experimental analyses and biochemical verification.

Thorough analyses of microsporidian genomes are essential to identify and functionally characterize species-specific proteins, which can provide novel drug targets to fight microsporidiosis in humans as well as environmentally and economically important animals. The identification of potential drug targets requires reliable tools to accurately identify and characterize divergent genes in microsporidia. Our approach improves the quality and quantity of functional genome annotation of a divergent organism and presents the first high-quality genome and annotation of the microsporidian *V. necatrix*.

Even though our approach requires a manual curation step, structural homology tools for protein annotation are an important complement to traditional sequence annotation tools and aid in overcoming annotation challenges with divergence and long evolutionary distances. We expect structural homology searches to become even more powerful as additional reference structures become available and as structural prediction tools continue to improve. Our plugin is a valuable tool for the accurate functional annotation and curation of genomes obtained from highly divergent, non-model organisms.

## Methods

### *V. necatrix* genomic DNA extraction

*V. necatrix* spores were propagated in the fourth and fifth instar larvae of Helicoverpa zea (corn earworm). The larvae were homogenized in Fisher 50 mL closed Tissue Grinder System tubes in water, filtered through a double layer of cheesecloth, and further filtered through 100 and 40 µm Biologix centrifugal filters before storage at –80 °C until further use. For genomic DNA extraction, *V. necatrix* spores were thawed, purified over 100% Percoll, and washed three times with sterile MilliQ water before the spore homogeneity was assessed by light microscopy. 12 mg of highly pure spores were germinated using the alkaline priming method (*82*). Spores were resuspended in 200 µl 0.1 M KOH for 20 minutes at 22 °C, pelleted via centrifugation at 2000 x g for 2 minutes, and resuspended in 100 µl germination buffer (0.17 M KCl, 1 mM Tris-HCl pH 8.0, 10 mM EDTA). A germination rate of approximately 80% was observed by light microscopy. To extract genomic DNA from the germinated spores, the Monarch® Genomic DNA Purification Kit (NEB, Cat# T3010) was used (10 µl Proteinase K, 3 µl RNase). Genomic DNA was eluted twice with 80 µl sterile MilliQ water. DNA quantification and qualification were assessed by Nanodrop and Qubit. Additional DNA quality assessments included electrophoresis on a 0.8% agarose gel stained with ethidium bromide and PCR amplification of a control gene.

### Sequencing and assembly

The extracted *V. necatrix* genomic DNA was sent to the National Genomics Infrastructure (NGI) Uppsala Genome Center (Science for Life Laboratory, Uppsala, Sweden) for PacBio *de novo* sequencing. To prepare the sequencing library for PacBio sequencing, 2 µg of genomic DNA were sheared on a Megaruptor3 instrument (Diagenode, Seraing, Belgium) to a fragment size of about 18 kb. The SMRTbell library was prepared according to PacBio’s Procedure & Checklist – Preparing HiFi Libraries from low DNA input using SMRTbell Express Template Prep Kit 2.0 (Pacific Biosciences, Menlo Park, CA, USA). The SMRTbells were sequenced on a Sequel II instrument, using the Sequel II sequencing plate 2.0, binding kit 2.2 on one Sequel® II SMRT® Cell 8M, with a movie time of 30 hours and a pre-extension time of 2 hours.

The sequencing resulted in 2’053’200 reads with a total of 28 gigabases and an N50 read length of 13.7 kb. The dataset was split into 14 read sets. Read sets were assembled using hifiasm 0.16.1. The resulting assemblies were split into 4 pseudo-haplotypes and telomere-to-telomere contigs were selected for each chromosome. The final assembly was then polished using Flye.

### Sample preparation for RNA seq

20 mg of highly germination-competent *V. necatrix* spores (>80% germination efficiency), stored at –80 °C, were thawed and cleaned by centrifugation through a 50% Percoll cushion. Subsequently, three MilliQ water washes were performed to remove Percoll remnants. Germination of cleaned spores was performed by alkaline priming of the spores in 200 µl of KOH followed by adding germination buffer (0.17 M KCl, 1 mM Tris-HCl pH 8.0, 10 mM EDTA). Gemination events were confirmed by light microscopy followed by the immediate addition of 300 µl of Ex-Cell 420 medium supplemented with 1 mM ATP. The sample was immediately added to an equal volume of Trizol reagent (Invitrogen Cat no. 15596026) and further supplemented with ⅓ volume of zirconium beads. Samples were vortexed for 1 minute and incubated on ice for 1 minute. This step was repeated two more times. Samples were spun down at 20,000 x g for 10 minutes at 4 °C followed by withdrawal of the aqueous layer and two subsequent extractions of the aqueous layer with chloroform. Overnight RNA precipitation was done with 2.2 volumes of ice-cold 96% ethanol, 1/10 volume of 3 M sodium acetate (pH 5.2), and 1 µl of Glycol blue co-precipitant. The next day, RNA precipitates were pelleted by centrifugation and washed twice with ice-cold 75% ethanol. The pellet was dissolved in 20 µl of nuclease-free water and treated with RNase-free DNase 1 (Invitrogen EN0521). As control and confirmation, the RNA sample was run on a 2% agarose gel.

### RNA Library Preparation and NovaSeq Sequencing

RNA samples were quantified using Qubit 4.0 Fluorometer (Invitrogen, Carlsbad, CA, USA), and RNA integrity was checked with an RNA Kit on an Agilent 5300 Fragment Analyzer (Agilent Technologies, Palo Alto, CA, USA). RNA sequencing libraries were prepared using the NEBNext Ultra RNA Library Prep Kit for Illumina following the manufacturer’s instructions (NEB, Ipswich, MA, USA). Briefly, mRNAs were first enriched with Oligo(dT) beads. Enriched mRNAs were fragmented for 15 minutes at 94 °C. First-strand and second-strand cDNAs were subsequently synthesized. cDNA fragments were end-repaired and adenylated at 3’ ends, and universal adapters were ligated to cDNA fragments, followed by index addition and library enrichment via limited-cycle PCR. Sequencing libraries were validated using the NGS Kit on the Agilent 5300 Fragment Analyzer (Agilent Technologies, Palo Alto, CA, USA), and quantified with the Qubit 4.0 Fluorometer (Invitrogen, Carlsbad, CA, USA).

### RNA seq data quality assessment

RNA seq data was received as fastq reads. Quality was checked with FastQC and sequences were subsequently subjected to trimming using Trimmomatic (v0.33) (https://github.com/timflutre/trimmomatic) to remove adapter contaminations (removal of first 120 bp) using the option “ILLUMINACLIP:TruSeq3-PE.fa:2:30:10:2:keepBothReads LEADING:3 TRAILING:3 MINLEN:120”. The trimmed reads were then aligned to the predicted proteome with STAR (v2.7.10). Of all aligned reads, 79.53% were uniquely mapped reads, 4.97% of reads mapped to multiple loci, 6.19% of reads mapped to too many loci, and 9.29% of reads were unmapped. Unmapped reads could be poor-quality reads, missed genes in the original gene prediction, or contamination from the host.

### Gene prediction and annotation

Prior to ORF prediction, potential transposable elements (TE) were identified using RepeatModeler, a de novo transposable element (TE) identification and modeling package. Using default parameters, a database of TE families was built. Next, RepeatMasker was used to softmask the genome followed by gene prediction with ProtHint and Augustus via the BRAKER pipeline. The quality of the predicted genes was assessed using BUSCO (v5.4.3) against the microsporidia_odb10 dataset.

### Generating a database for the functional annotation with our ChimeraX annotator plugin

For the functional annotation, a database was generated to retrieve the best sequence and structure-based matches for each input sequence. The sequence-based search was done using Diamond with the ultra-sensitive option against the non-redundant NCBI database. The eggNOG mapper allowed for functional annotation based on orthology predictions which is considered more precise than traditional homology searches. For structural matches we folded the *V. necatrix* proteome and the hypothetical proteins of *E. cuniculi* using ColabFold with default parameters. Next, we used each individual predicted 3D structure as input for Foldseek searches employing the alignment type 3Di+AA Gotoh-Smith-Waterman (local, default) and ran it against three different databases: 1. PDB, 2. AlphaFold database from the 20 first annotated model organisms (accession date: 07-15-2022), one representative of each microsporidian clade (**Figure 1a**), and 3. SwissProt AlphaFold. Additionally, for the *E. cuniculi* proteins, individual, well-predicted protein domains were automatically separated using the Predicted Aligned Error (PAE) (*38*) and subjected to the TM-align algorithm in Foldseek. As a measure of confidence, the E-value is displayed for all Diamond and eggNOG searches, while the significance of the Foldseek searches varies with the alignment type: The bit score assesses 3Di+AA Gotoh-Smith-Waterman search results and the TM score (global score) represents the confidence of TM-align searches. Further, to predict the overall 3D structure and the presence of an SP or TMD for each analyzed protein, the Deep Transmembrane Helix Hidden Markov Model (DeepTMHMM, v1.0.20) software was used.

To combine and display the generated information and homology matches for each *V. necatrix* input sequence, we developed a ChimeraX annotator plugin (**Supplementary Figure 2**). It retrieves a list of all predicted *V. necatrix* protein 3D structures, shows the eggNOG annotation in the user interface (**Supplementary Figure 2b**), and presents a list of structural matches and sequence-based hits, respectively, along with corresponding confidence values. Further, the proteins corresponding to structural hits can be superimposed with the *V. necatrix* protein of interest, allowing for visual inspection of the structure match. Additionally, the overview of all structural and sequence hits per protein allows for manual curation and functional annotation according to the best match.

### Installation and Use of the Plugin

After successful installation of the plugin, using command “devel build PATH_TO_ANNOTATER_FOLDER; devel install PATH_TO_ANNOTATER_FOLDER; devel clean PATH_TO_ANNOTATER_FOLDER” in the ChimeraX command line, the user can load the previously created proteome predictions, by selecting the “Data folder” path and pressing the “Search” button (**Supplementary Figure 2b**). The structure prediction for the first protein-coding gene will be displayed in the ChimeraX viewer, and the tool window will show the gene ID, a potential eggNOG annotation with a description of the protein function and E-value, if applicable, and a DeepTMHMM prediction that can be a global protein (GLOB), a TMD, a SP, or both TMD and SP. Below these descriptions, the structural matches from PDB (blue), AlphaFold microsporidian proteomes, and the available 20 model organisms (red) and AlphaFold Swissprot (yellow) are listed followed by the Diamond sequence-based blast hits (**Supplementary Figure 2b**). All entries are sorted according to their assigned E-value which can help to assess the certainty of a structural or sequence-based match.

The protein of interest is displayed as a complete model #1 in rainbow colors according to the AlphaFold confidence (**Supplementary Figure 2a, d left panel**). If the domains’ position relative to each other is of high uncertainty, the model can be hidden in the Models window (**Supplementary Figure 2a, lower panel**) and the individual, well-predicted domains become visible. Further, single domains in the list can have a subcategory that indicates a “missing structure” due to the exclusion/removal of highly disordered regions or flexible linkers. To visually inspect the structural matches, the user can click on an entry and subsequently, the corresponding protein structure will be superimposed (automatically applying the matchmaker in ChimeraX) to the investigated protein in the model viewer (**Supplementary Figure 2a, d right panel**). The superimposed structure can either be a single protein or a protein complex. In the latter case, the protein of the complex with structural homology to the protein of interest will be highlighted in green. If the structural match is from a PDB entry, the protein “chain” of the complex matching the protein of interest is added as a suffix to the PDB ID as shown in **Supplementary Figure 2a** (PDB 4pz6_A) and **d** (PDB 3j96_G). Further, for any selected structural hit an info panel will appear in the console log listing “ID”, “Title”, “Chain information” and “Parameters” (**Supplementary Figure 2c**). Once the user has found an adequate functional annotation, enabling the checkbox “Overwrite annotation on click” allows the protein being investigated to be named after the matching protein by clicking on the corresponding entry (**Supplementary Figure 2b**). The complete genome annotation can be saved in tsv-format facilitating accessibility of the file and further editing in a suitable program. Our ChimeraX-Plugin was successfully tested using ChimeraX version 1.3 and 1.4.

### Analysis of false positive ORF prediction of non-annotated genes

To estimate how many of the predicted hypothetical genes might be false positive ORFs, we compared the RNA sequencing reads between annotated and non-annotated genes (**Supplementary Figure 4a**). More than 87% of the hypothetical genes are covered by RNA reads, which is close to the 92% coverage of the successfully annotated genes and suggests that most hypothetical genes are present. The hypothetical ORFs could either encode yet unknown proteins or are the result of an overestimated number of protein-coding regions predicted by BRAKER. However, more than 550 of these ORFs have an mRNA sequence count over 200 (**Supplementary Figure 4a**). Further, a significantly higher number of proteins with an SP and TMD is predicted among the hypothetical compared to the classified proteins (**Supplementary Figure 4b**). Since both SPs and TMDs seem to be key features of host-exposed proteins (*74*), this enrichment suggests that many of the hypothetical proteins belong to the group of exported proteins. Host-exposed proteins have been found to evolve faster than the remainder of the proteome, presumably because these proteins are under pressure from the host immune system. In the Nematocida it was shown that host-exposed proteins evolve rapidly and are most often lineage-specific (*74*). The majority of these proteins are thus hypotheticals and present a low evolutionary traceability which hinders further annotation efforts.

### Benchmarking

Shortly after we completed the functional genome annotation, the automated annotation tool ProtNLM was published and represented the new standard for sequence-based annotation, replacing eggNOG. Therefore, we decided to benchmark our approach, and we manually compared the final gene function annotations that we generated with our method for the *V. necatrix* genome and the *E. cuniculi* (strain GB-M1) uncharacterized proteins from UniProt (*8*) to the results from ProtNLM. We distinguished between identical annotations, different annotations, not-identified annotations, and experimentally determined gene functions which are based on published studies. The identical annotations also include cases where either tool, ours or ProtNLM, predicted a gene function and the respective other tool predicted only a protein domain that is typically involved in this gene function. Differing predictions also include a subsection of potential miss-annotations made by ProtNLM. Not identified gene functions comprise hypothetical proteins, uncharacterized proteins, and DUF-domain-containing proteins, independently of whether a feature like TMD or SP is listed. Additionally, for the *E. cuniculi* uncharacterized gene set, we differentiated between the characterized protein and the characterized protein domain. The number of proteins in each category was counted and displayed in a pie chart to visualize the performance of our annotation approach.

### RBL identification, analysis, and visualization

To identify RBL proteins in the order Nosematida using structural homology, characterized RBL domain-containing proteins, such as PTP4 and PTP5, were identified in *V. necatrix* and *E. cuniculi*. The corresponding AlphaFold models were extracted from the annotator database and large disordered regions were trimmed to retain only the ricin-type β-trefoil lectin domain. This domain served as a template for structural homology searches in Foldseek using the TM-align algorithm. Among the homology matches were mannosyl transferases, which typically contain a functional RBL domain (i.e., VNE69_06039 and VNE69_12061) and were thus removed. HMMER profiles (v3.3.2) (http://hmmer.org) (*83*) were generated to detect RBL proteins that were potentially overlooked by the structural search.

Next, the sequences of all identified RBL domain-containing proteins were aligned with MUSCLE (v5) (*84*) and trimmed with trimAI (*85*), a tool for the automated removal of spurious sequences and poorly aligned regions from a multiple sequence alignment. The remaining sequences were used to build a cladogram with IQTREE2 (*86, 87*) with 1000 bootstrap replicates and the MFP option for choice of substitution model.

To generate a structural network graph, the visualization software Gephi (*88*) was used according to the user guidelines. Briefly, the required nodes and edges data sheets were generated for which the squared TM score served as edge weight. Data was imported into Gephi, the graph type was set to undirected, statistic tools In/Out Degree, Network Diameter, Graph Density, Modularity, and Average Clustering were run using default settings, and ForceAtlas 2 was chosen as layout.

## Declarations

### Availability of data and materials

The raw PacBio data and the final *V. necatrix* assembly including annotation information were deposited at NCBI under BioProject accession ID PRJNA909071 and BioSample SAMN32066506. Raw transcriptomics data were deposited under PRJNA909071. The ChimeraX plugin and the databases built for annotation of the *V. necatrix* and the *E. cuniculi* genes can be accessed on Zenodo under 10.5281/zenodo.7974739 (https://doi.org/10.5281/zenodo.7974739).

### Authors’ contributions

J.B., together with D.S., R.R.W., A.B., and C.R.V. conceived the study. D.S. wrote all scripts and performed the computational work, and together with R.W. and A.B. performed the annotation work. R.R.W. cultivated microsporidia, extracted the genomic DNA, and H.S. performed the RNA sequencing-related experimental work. All authors interpreted the results, and wrote, and edited the manuscript.

## Acknowledgments

We thank all members of the Barandun laboratory for the helpful discussions. The computations were enabled by resources provided by the Swedish National Infrastructure for Computing (SNIC) at High-Performance Computing Center North (Project Nr. SNIC 2021/23-718 and SNIC 2021/22-936), partially funded by the Swedish Research Council through grant agreement no. 2018-05973. The authors acknowledge the support of the National Genomics Infrastructure (NGI) / Uppsala Genome Center and UPPMAX for assisting in massive parallel sequencing and computational infrastructure. Work performed at NGI / Uppsala Genome Center has been funded by RFI / VR and Science for Life Laboratory, Sweden. Further, support by NBIS (National Bioinformatics Infrastructure Sweden) is gratefully acknowledged.

