## Supplementary Information for "Functional annotation of a divergent genome using sequence and structure-based homology"

**Supplementary Figure 1. Complementary annotation pipeline from genome to function.**

**Supplementary Figure 2. ChimeraX Annotator plugin overview.**

**Supplementary Figure 3. Structural network of ricin B lectins in Nosematida shown with organism and gene IDs.**

**Supplementary Figure 4. RNA sequencing reads of annotated and unannotated genes, and protein features of hypothetical, uncharacterized, and classified proteins.**

**Supplementary Table 1. Annotation table *V. necatrix*.** Table of all locus tags and the resulting structure-based annotations. Further, the ProtNLM results are included and the differences to our annotation are indicated (x: different annotations, y: non-identified, z: potential mids-annotations, e: structural). In addition, the spore-0hr RNA sequence counts are listed per gene.

**Supplementary Table 2. Annotation Table *E. cuniculi.*** List of 381 uncharacterized proteins from E. cuniculi and the updated annotation using our approach.

**Supplementary Data File 1. Data file with the ChimeraX plugin, the used annotation database and all predicted structures.** DOI 10.5281/zenodo.7974739 https://doi.org/10.5281/zenodo.7974739) V_necatrix_alphafold.zip (Alphafold models and associated files for all *V. necatrix* proteins), chimerax_annotater_plugin.zip (ChimeraX plugin install file), v_necatrix_annotation_data.zip (annotation data used in the ChimeraX plugin and generated as described in the methods section)

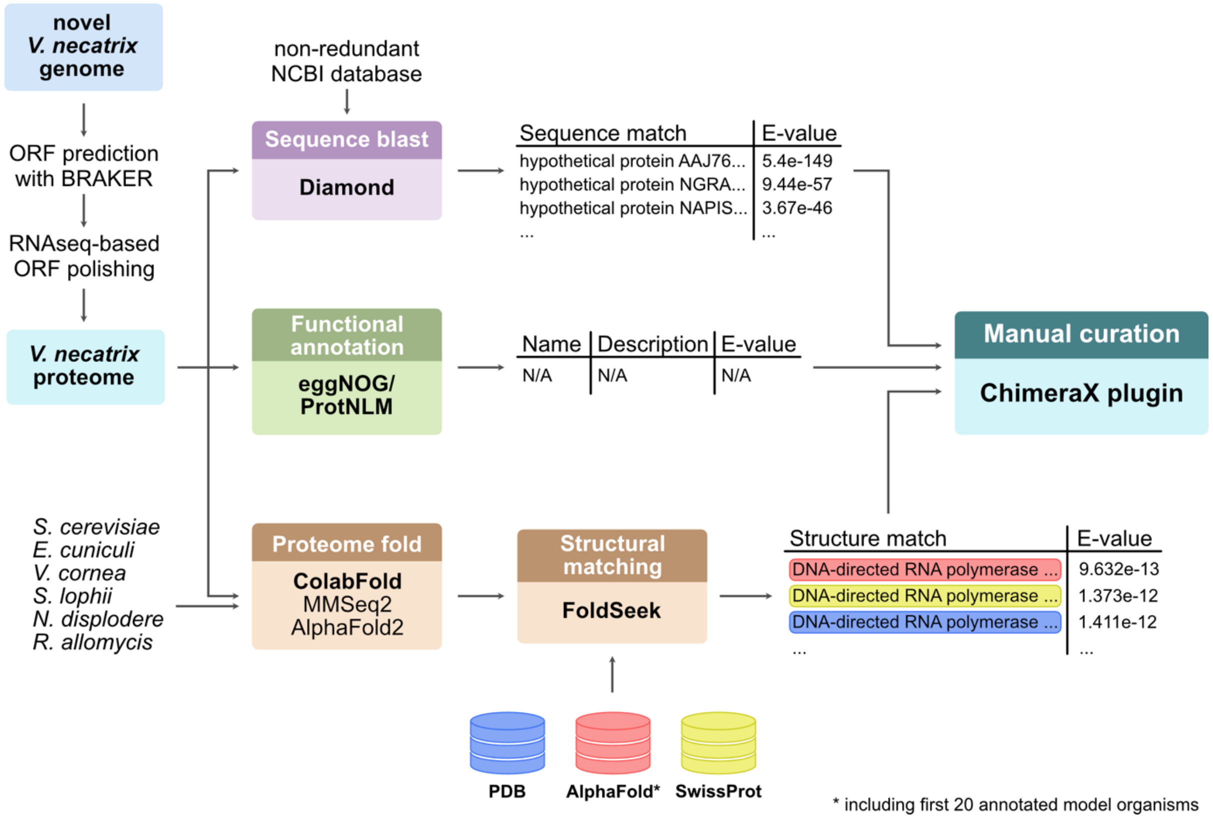


**Supplementary Figure 1. Complementary annotation pipeline from genome to function.** We predicted ORFs in our novel *V. necatrix* genome using BRAKER and polished the protein-coding regions with our transcriptomic data. The corresponding *V. necatrix* proteome served as basis for functional annotation using a combination of three different approaches: Sequence-based annotation with Diamond, functional annotation via domain homology using eggNOG and via natural language processing based on the amino acid sequence operated by ProtNLM, and structural homology searches employing ColabFold and Foldseek. For the structural similarity search, we used ColabFold to fold the proteomes of *V. necatrix*, *S. cerevisiae* as representative model organisms, and five microsporidian species, each representing a clade. Among the folded proteomes, we searched for structural homologs to the *V. necatrix* proteins and further used the databases PDB, AlphaFold/Proteome (accessed July 2022, only 20 folded proteomes of model organisms were available) and AlphaFold/SwissProt. The functional prediction matches and the corresponding E-value, bit score, or TM score of all three approaches were visually combined in our ChimeraX annotator plugin allowing us to find the best matches and manually curate the functional annotations.

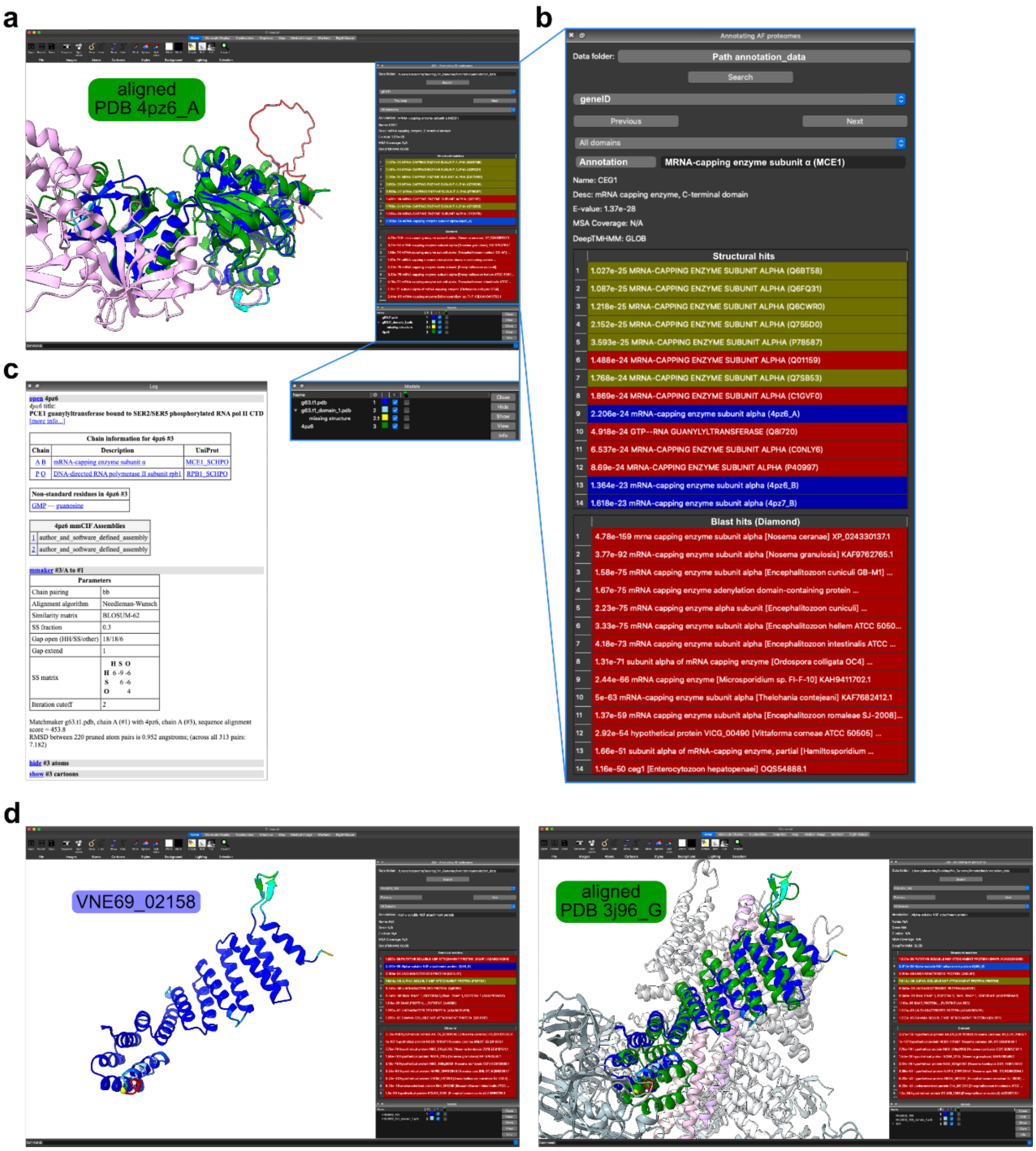


**Supplementary Figure 2. ChimeraX Annotator plugin overview.** (**a**) ChimeraX window and plugin showing the AlphaFold-predicted protein encoded by gene 63.t1 (now renamed VNE69_03064) and superimposed with a structural match (PDBid: 4pz6_A) in the 3D viewer. Lower panel: Zoom of Model window listing structures, domains, and missing structures displayed in the model viewer. (**b**) Our ChimeraX plugin "JSB-Annotating AF proteomes" consists of a tool window docked to the right, from top to bottom: "Data folder" path search tool, gene ID bar, domain selection bar, potential eggNOG annotation with protein name, description and E-value, MSA coverage, DeepTMHMM information, a list of structural matches from databases PDB, AlphaFold microsporidian proteomes, and AlphaFold SwissProt, followed by Diamond sequence blast hits. (**c**) ChimeraX Log console showing name, chain information, and parameters of the selected PDB structure. (**d**) ChimeraX with the plugin presenting the AlphaFold protein structure encoded by gene VNE69_02158 in rainbow colors (left panel) and superimposition with a structural match (PDBid: 3j96_G) in green (right panel).

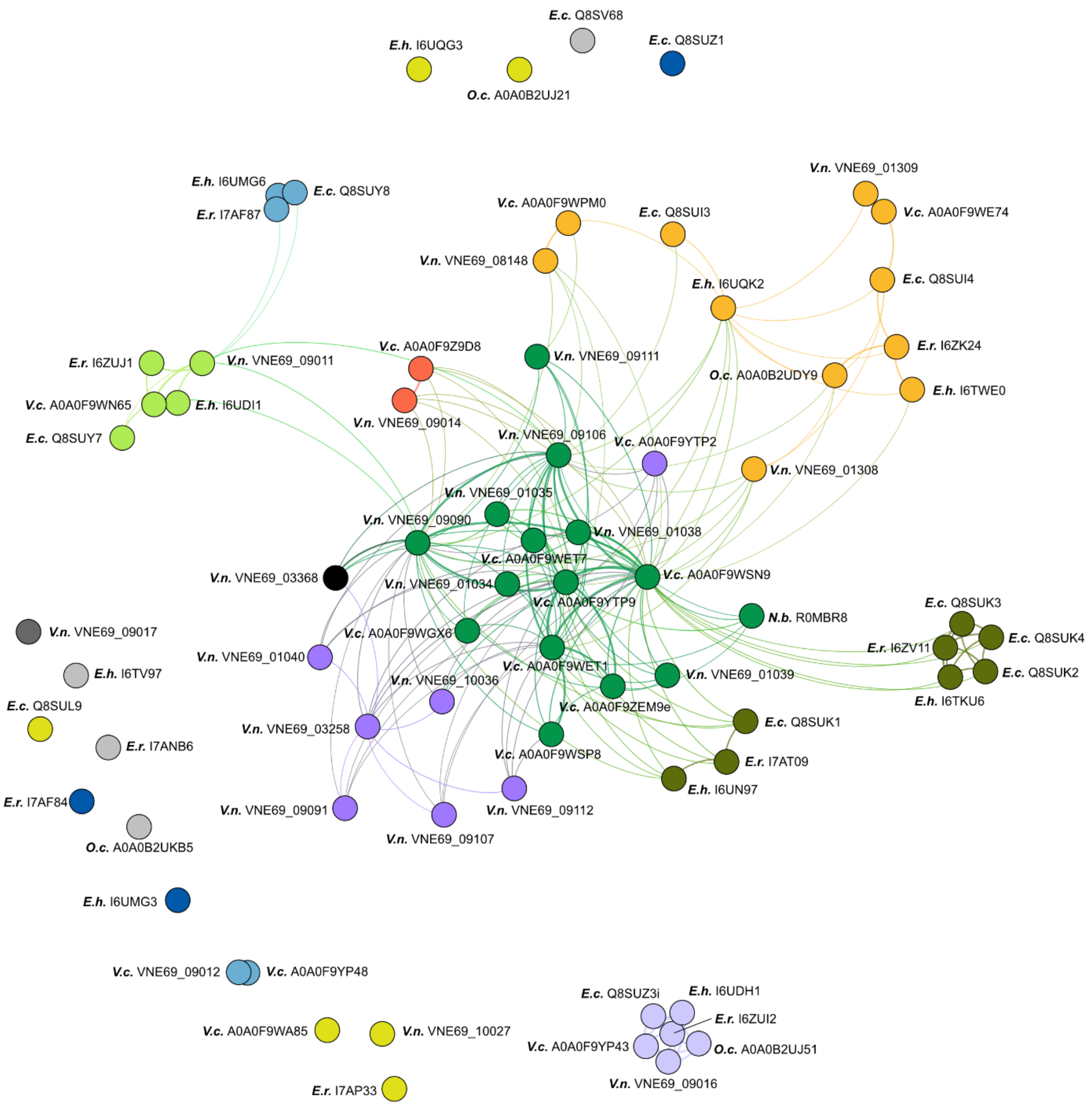


**Supplementary Figure 3. Structural network of ricin B lectins in Nosematida shown with organism and gene IDs.** Protein folds of all Nosematida RBLs identified in this study were predicted with AlphaFold and RBLs were clustered according to structural similarity based on their TM score using Gephi. RBLs are color-coded according to their clade in **Figure 4a**. Each node represents one RBL protein, and connecting lines indicate the degree of structural relatedness.

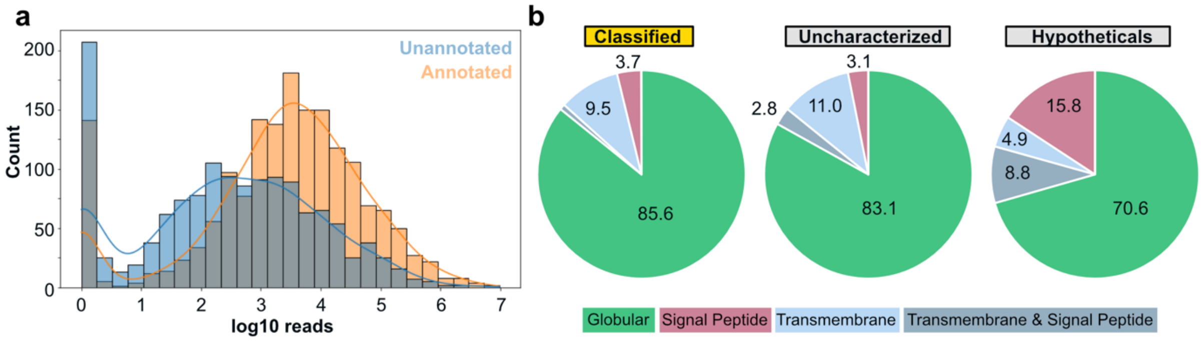


**Supplementary Figure 4. RNA sequencing reads of annotated and unannotated genes, and protein features of hypothetical, uncharacterized, and classified proteins. (a)** Distribution plot for RNA sequencing reads of annotated vs. unannotated genes. The bar plot is non-stacked, and the blue or grey bars correspond to unannotated genes and the orange bars correspond to annotated genes. (**b**) Pie plots of the classified, uncharacterized, and hypothetical gene groups with the percentage of globular (green), signal-peptide-containing (coral), transmembrane-containing proteins (light blue), and those with both TMD and SP (grey blue).
